# Basal Ganglia Control of Reflexive Saccades: A Computational Model Integrating Physiology Anatomy and Behaviour

**DOI:** 10.1101/135251

**Authors:** Alex J. Cope, Jonathan M. Chambers, Tony J. Prescott, Kevin N. Gurney

## Abstract

It is hypothesised that the basal ganglia play a key role in solving the problem of action selection. Here we investigate this hypothesis through computational modelling of the primate saccadic oculomotor system. This system is an excellent target for computational modelling because it is supported by a reasonably well understood functional anatomy, has limited degrees of freedom, and there is a wealth of behavioural and electrophysiological data for model comparison. Here, we describe a computational model of the reflexive saccadic oculomotor system incorporating the basal ganglia, key structures in motor control and competition between possible actions. To restrict the likelihood of overfitting the model it is structured and parameterised by the known anatomy and neurophysiology along with data from a single experimental behavioural paradigm, then validated by testing against several additional behavioural experimental data without modification of the parameters. With this model we reproduce a range of fundamental reflexive saccadic results both qualitatively and quantitatively, comprising: the distribution of saccadic latencies; the effect of eccentricity, luminance and fixation-target interactions on saccadic latencies; and the effect of competing targets on saccadic endpoint. By investigating the model dynamics we are able to provide mechanistic explanations for the sources of these behaviours. Further, because of its accesibility, the oculomotor system has also been used to study general principle of sensorimotor control. We interpret the ability of the basal ganglia to successfully control saccade selection in our model as further evidence for the action selection hypothesis.

## 1 Introduction

Saccades are ballistic eye movements that direct visual attention to putative targets of interest. Given the primacy of vision as a sensory modality in humans, saccades are clearly essential for our goal-directed behaviour. Reflexive saccades are elicited by the phasic appearance of targets peripheral to the current fixation, and are to be contrasted with voluntary saccades which are under cognitive control via, say, verbal instruction or memory recall [1].

The selection of targets for saccade generation in the oculomotor system is a good target for computational modelling for several reasons. First, the behavioural output may be characterised with only three degrees of freedom (dof), of which we are often only interested in two (for horizontal and vertical saccades). This is to be contrasted with upper limb movement, say, which has seven dof (or many more if we work at level of the musculature) [2]. Second, there is a wealth of behavioural data characterising saccades - see for example, [3–6] - which has simple interpretation because of the 2-dof constraint. Often this data takes the form of reaction time and/or saccade location to stimulus onset in simple paradigms with one or two targets. Third, the anatomy of the neural substrate is particularly well understood [7, 8] offering the opportunity for modelling with biologically plausible architectures. Finally, there is a corresponding abundance of electrophysiological data from single unit recordings in awake behaving monkeys [9] which facilitates model constraint and validation.

The basal ganglia (BG) have been implicated in the ‘action selection’ problem [10,11]: how out of several competing possibilities for action an animal cleanly chooses the best option. Holistically the BG sits in the centre of the oculomotor system, receiving input directly from the terminal retinatopic regions of the cortex, the Frontal Eye Fields (FEF), and indirectly from the retinatopic brainstem regions of the superior colliculus. The BG output inhibits circuits to shape the buildup of neural activity in both of these regions. The BG is clearly positioned to act as a central switch for action selection in the oculomotor system. However, superficially it may seem that this problem is divorced from simple reflexive saccade tasks, where often there is only a single target and thus a single option for action. However, electrophysiological studies have shown activity patterns in the BG during such tasks that imply a role for the BG in their performance. Despite extensive modelling work focusing on this part of the brain (Girard REFS) the influence of the BG on reflexive saccades has not been explored. Here we investigate this relationship through computational modelling.

In our previous work we described a simple computational model of the BG as an action selection system in isolation [11,12], and this work has been used as the basis of more extensive models of oculomotor function [13] exploring predominantly voluntary saccadic tasks that require cognitive competencies. Here we instead focus on the role of the BG in purely reflexive saccades, where the eye moves to fixate targets with abrupt onsets. This reduced scope allows us to create a model without prefrontal cortical regions, instead focusing on the better characterised sensory cortical and subcortical areas. One important principle in building our model is to facilitate validation of the model. This is undertaken using a two stage process, with the experimental evidence divided into two sections - one for fitting the model, after which all parameters are fixed - and one for testing the model without alteration of the parameters of the model, but instead simply providing different visual inputs. The first section of evidence includes anatomical and electrophysiological evidence, along with a single behavioural task. The second section contains further behavioural tasks. It would not be expected that the model would reproduce the second section of data using parameters derived from the first, and this therefore provides a strong test of the validity of the model, contingent upon the tasks being suitably distinct between the sections. The relationship of the model presented here to other models in the literature will be explored in the Discussion.

## 2 Methods

### 2.1 Model overview

In the Introduction we outlined that the modelling process we undertook contained two stages. Our model demonstrates reflexive saccades, and as such we are replicating data with many repetitions in which there is no learning, or any learning will have reached a steady state. Therefore we have no learning processes in the model, and the structure and parameters of the model are not subject to change once they have been set. Thus, we have a first stage of modelling in which we describe the structure of the model based upon the neuroanatomy and functional roles of the brain regions involved in reflexive saccades, and set parameters based on neurophysiological data, along with a single behavioural experiment. These parameters are then fixed for the second stage of the modelling process, where we validate the performance of the model against additional behavioural experiments using the parameters derived soley from the first stage.

The functional architecture of our model is constrained by the known anatomy and physiology of the oculomotor system [14] and includes: basal ganglia (BG), thalamus, superior colliculus (SC) and frontal eye fields (FEF). The macroscopic connectivity is shown in Fig 1. The form of the architecture is close to that described by N’Guyen et al [13].

**Fig 1.**
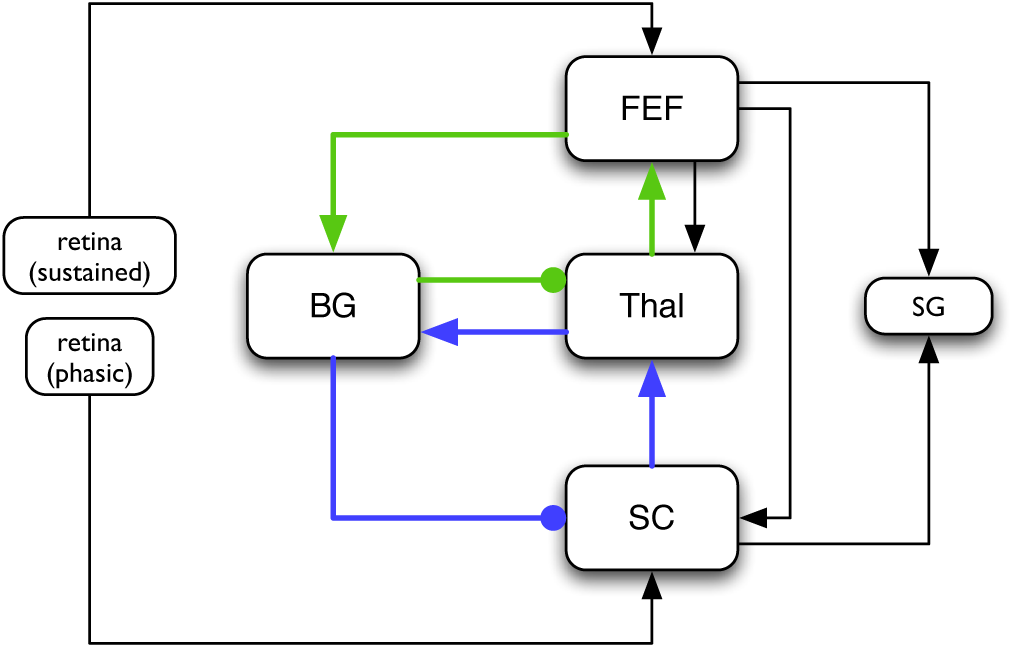
The macroscopic architecture of the model. The main nuclei modelled as brain systems are: basal ganglia (BG), frontal eye fields (FEF), thalamus (Thal), and superior colliculus (SC). Other components include the retina (providing sustained and phasic input) and the saccadic generator (SG) (motor output) which are not modelled in a biologically plausible way. The loops through basal ganglia, defining the architecture, are shown by coloured lines: the cortical loop (through FEF and Thal) in green, and the sub-cortical loop (through SC and Thal) in blue. Connections with arrowheads/filled circles indicate excitatory/inhibitory connections respectively.

As noted in the Introduction, our perspective here focuses on the basal ganglia and their role in action selection. The basal ganglia are connected in closed looped circuits with cortex, via thalamus [15–17], and also form loops with sub-cortical circuits [18]. Their outputs are tonically active and inhibitory, and selection is achieved by selectively releasing inhibition on cortico-thalamic or subcortical targets that encode specific actions [19].

Under the action selection hypothesis developed by Redgrave et al. (1999), each action is encoded in neural representations throughout a loop with basal ganglia; we refer to the anatomical substrate for each action encoded in this way as an *action channel,* or simply a ‘channel’. The strength of the input into these channels is termed the *salience* of the action represented by that channel. A high salience indicates that an action is more behaviourally relevant in the current context, and a low salience indicates that an action is less behaviourally relevant. In simple models of basal ganglia and cortico-thalamic loops, these channels comprise discrete, non-overlapping circuits [11,12,20]. We will have need to finesses this scheme in the current model, as described in section 2.2. Nevertheless, in broad terms, release of inhibition from basal ganglia to the thalamo-cortical or subcortical target in a channel allows activity in the target to build up and eventually reach a threshold, thereby allowing behavioural expression of the corresponding action.

In the oculomotor circuit, there are two principal loops through basal ganglia: a cortical loop originating in FEF [17], and a subcortical loop originating in SC [18]: see Fig 1. The FEF and SC are described in more detail below, but in the meantime, we note that, their respective loops with basal ganglia interact within a complex of thalamic nuclei, linking the FEF, SC, and the basal ganglia [21]. In addition the FEF innervates SC [22] and both FEF and SC control the saccadic generator (SG) in the brainstem which operate the musculature surrounding the eye to produce saccades [23].

Several regions of the brain that are associated with eye movements have been omitted from this model. Thus, the early visual processing stream in cortex, from V1 through to the lateral intraparietal region (LIP) (which then innervates FEF), has been subsumed into a ‘sustained retinal’ signal. Our rationale here is that we are interested in phenomena surrounding saccades to simple luminance targets which do not require the detailed feature extraction and analysis performed by these visual areas. Another region of cortex that is closely linked with eye movements is the supplementary eye fields (SEF) [24]. However, the SEF are involved only in the programming of sequences of saccades [25] and memory guided saccades [26–28]. Visually guided saccades, which are our remit here, are unaffected by lesion of SEF [29] and so we therefore omit them.

At the neuronal level, the model uses rate-coded *leaky integrator* units to represent small populations of neurons [30]. In brief, each unit uses a weighted sum of its inputs as the forcing term in a first order dynamical equation with a state variable we call the *activation a*,

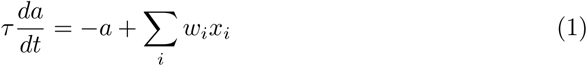

where *τ* is a time constant. This activation is used, in turn, as the input to a piecewise linear transform with saturation to form the final output, thereby constraining the output between 0 and 1,

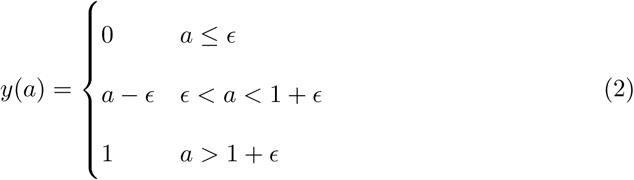

Here, *ϵ* defines an offset to simulate (if positive) the effect of tonic activity or (if negative) a threshold above which activity must rise for there to be an output. The exception to this rule is the STN which has an exponential output function which better approximates its physiology [31]:

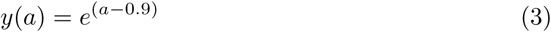

The model as a whole comprises assemblies of these units in each nucleus. Each assembly is referred to as a ‘layer’ since, in all cases, there is a retinatopically defined spatial organisation defined by a 50-by-50 unit array (see Fig 3). Connections between layers comprise a variety of topographic connection schemes, and can be normalised by dividing each weight by the sum of all weights in the connection. We now turn to a more detailed description of component sub-systems, starting with the most complex – the basal ganglia.

### 2.2 Basal Ganglia

Our instantiation of basal ganglia is based around our previously published model (Fig 2)

**Fig 2.**
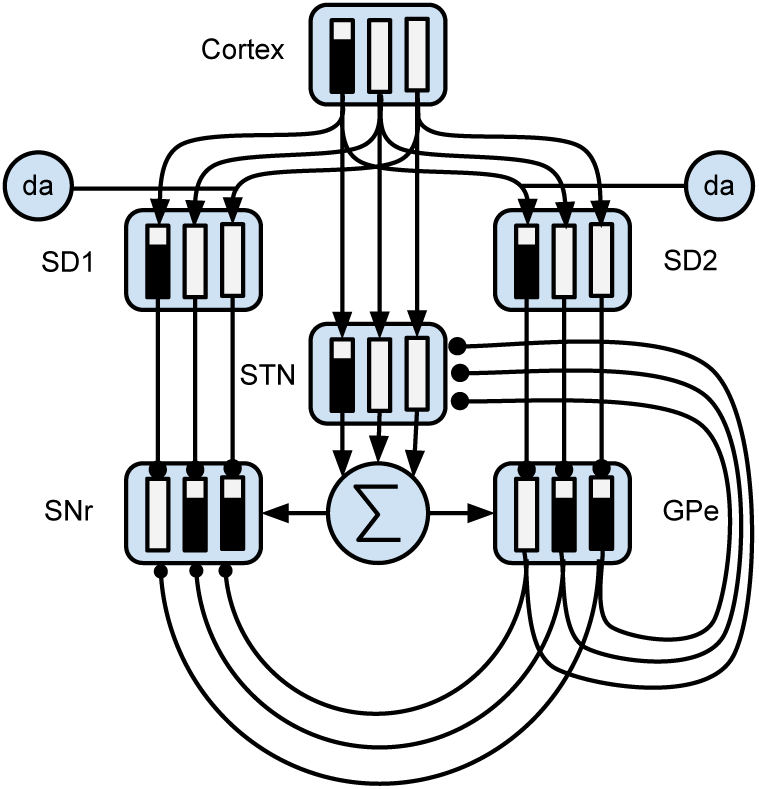
The basal ganglia model component. Connections with arrowheads/filled circles indicate excitatory/inhibitory connections respectively. The circles labeled ‘da’ provide dopaminergic influence on the inputs to the striatum (SD1 and SD2). Three action channels are shown to illustrate the selection mechanism. The small bar-graphs within each nucleus indicate the level of activity within each channel therein. For example, the channel indicated by the leftmost bar has a high salience (cortical input). The diffuse projection from STN is indicated by the summation symbol (diffuse projection is equivalent to summing channel-wise projections and dispersing focally

[11,12] which will be henceforth referred to as the *GPR model*. This model incorporates the following main components of the primate BG [32,33]: (i) The striatum – the main input station to the BG – which is divided into two interdigitated populations of projection neurons expressing primarily D1 or D2 type dopaminergic receptors (termed *SD1* and *SD2* respectively here); (ii) The subthalamic Nucleus (STN); (iii) the external segment of the globus pallidus (GPe); (iv) the output nucleus relevant for saccadic control – the substantia nigra pars reticulata (SNr) [14]

The connectivity of the GPR model (Fig 2) is constrained by the known anatomy and physiology of the BG [34]. Physiologically, the only source of glutamate within BG is the STN whose projections are therefore excitatory; all other nuclei have GABAergic projection neurons and are therefore inhibitory. The cortex sends glutamatergic projections to both the SD1 and SD2 populations in striatum which, in turn, project preferentially to the SNr, and GPe respectively [35]. The cortex also projects to the STN which sends diffuse projections to the SNr and GPe [36]. The GPe, in turn, projects to the SNr and to the STN.

Within BG, there are several mechanisms supporting competitive processing for selecting channels whose inhibitory output is reduced. The selection mechanism of the GPR model is the ‘off-centre, on-surround’ scheme proposed by Mink and Thach (1993). Here, the STN provides diffuse excitation (the ‘on-surround’) to the SNr, and the SD1 neurons in striatum provide focussed inhibition (the ‘off-centre’) to SNr. This arrangement leads to selection behaviour via release of target inhibition, since channels that have strong salience (input) have weak output at the level of SNr, and channels with weak salience have enhanced output.

The GPe is not included in the centre-surround circuit described above but still plays an key role in selection. Thus, the diffuse projection from STN can, if unchecked, cause massive excitation within each channel of SNr (since each one receives input from all STN channels). Gurney et al [11,12] showed that the inhibitory feedback from GPe to STN acts as an ‘automatic gain control’ to help prevent this.

At the neuronal level, the STN, GPe and SNr have tonic output levels [19,38,39]. This is modelled using piecewise linear output functions with positive offsets *∊* (see equation 2). In striatum, SD1 and SD2 have negative offsets mimicking the so-called ‘down-state’ of medium spiny neurons which have a resting potential far below spiking threshold and require massive co-ordinated input to generate action potentials [40]. In addition the SD1 and SD2 neurons are influenced by dopamine in different ways. Our model incorporates the neuromodulatory effects of dopamine, which has a predominantly facilitatory effect on cortico-striatal transmission at medium spiny neurons with D1 receptors [41,42] and an inhibitory effect on those with D2 receptors [43]. This is modelled by multiplying the cortico-striatal weights by factors of (1 + *d*) and (1 − *d*) for SD1 and SD2 respectively, where *d* represents the level of tonic dopamine.

The original GPR model had only six channels, with one leaky integrator unit per channel in each nucleus, and simple connectivity in which focussed inhibition from striatum to SNr and GPe was defined by a simple one-to-one scheme (with a connection from each unit in SD1/SD2 to a counterpart in SNr/GPe). In the oculomotor model, we made two modifications to this scheme to incorporate the continuous, topographic (retinatopic) connectivity required. First, each nucleus is comprised of a layer of leaky integrator units arranged into a two-dimensional grid of 50 by 50. Here, each unit represents the activity associated with a corresponding spatial location in the visual field. Second, the connections from striatum to SNr and GPe were defined by projective fields with many weighted connections. Specifically, each unit in SD1 projected to a counterpart SNr_*j*_ in SNr with some weight *w*_*max*_, but also connected to neighbouring nodes in SNr with a weight given by *w*_*max*_*G*(*d*), where *G*(*d*) is a circularly symmetric, 2D-Gaussian which is a function of distance *d* from SNr_*j*_ (see Fig 3). A similar scheme applied with respect to SD2 and GPe. In addition the strength of the excitatory outputs from the STN were decreased in comparison with their GPR counterparts.

**Fig 3.**
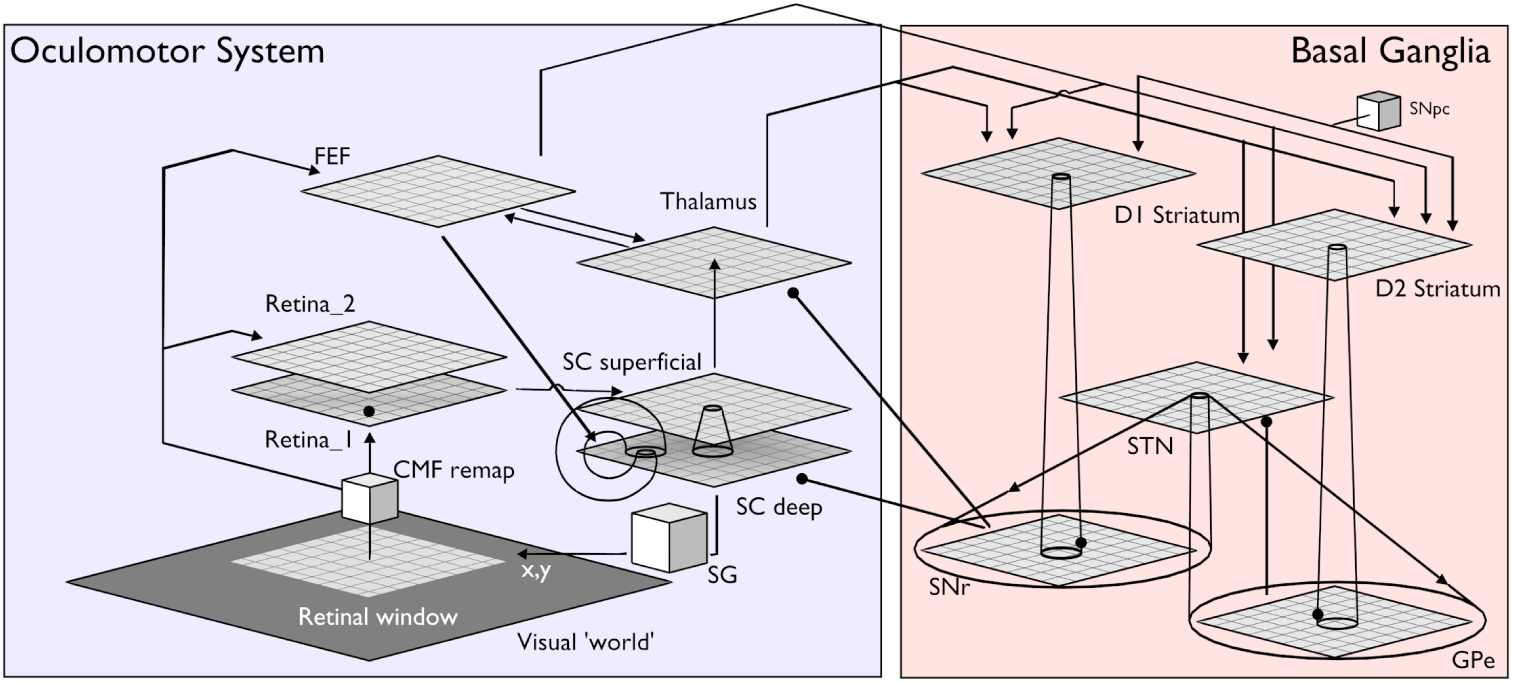
The detailed architecture of the model. Connections with arrowheads/filled circles indicate excitatory/inhibitory connections respectively. Conic section between layers represent spatial pattersn of connectivity weights governed by 2D Gaussians, and cones covering the entire target layer represent all-to-all connection patterns. Cubes represent components that are not implemented in a biological manner. The retinal layers 1 and 2 correspond to the ‘faster’ and ‘slower’ reacting layers respectively. They receive an input from a window on the world array, the location of which is determined by the current eye co-ordinates from the saccadic generator (SG). All other abbreviations are described in the text.

### 2.3 Loops through basal ganglia with frontal eye fields and superior colliculus

The Frontal Eye Fields (FEF) are a key cortical area for the generation of saccadic eye movements [14,25,44,45]. Saccadic targets are mapped retinatopically over the surface of FEF [44–46], and increased neural activity at a location in the map precedes a saccade to that location. Importantly, the FEF is also associated with visual decision making [47–50]. Thus, in a saccade choice task, increased FEF activity is predictive of the eye movement whether correct or incorrect [51], rather than the correct response. Remarkably, this prediction is reliable with a sampling of fewer than 7 neurons or trials [52].

FEF neurons can be divided into three functional groups, related to whether their activity corresponds with visual stimuli, motor action, or both [22]. Here we simplify this categorisation using a single layer of 50 by 50 units representing the mean of all three groups. This layer therefore responds to both visual stimuli and the buildup of activity associated with motor (saccadic) action. The retina provides a persistent luminance signal into the FEF through the dorsal visual pathway [53] which is abbreviated in our model to a direct connection with delay.

The FEF provides input into the BG [54] (at SD1, SD2, STN - see Fig 3) which, in turn, projects back to thalamus in a retinatopically organised way, [17,55]. In addition the thalamic targets of this path are regions with strong reciprocal connections to the FEF [56]. In this way the FEF forms channel-based loops through basal ganglia of the kind described above. Such circuits formed the basis the model of Humphries and Gurney [20]. In that model, the thalamic component included the thalamic reticular nucleus which implemented an additional selection mechanism (via a feed-forward competitive network within the cortico-thalamic loop). However, in the interests of model simplicity, this was not implemented in our oculomotor architecture.

The multiplicity of feedback loops through thalamus, cortex and basal ganglia can develop complex behaviour. In particular the thalamo-cortical loop may be thought of as an integrator, or accumulator of information whose ‘gain’ is modulated by inhibition from basal ganglia [57,58]. To understand how this modulation occurs mechanistically, we note two things: first evidence that projections from SNr form synapses proximal to the soma at thalamic neurons [59]; second, that proximal inhibition often acts to multiplicatively gate (and *in extremis* veto) excitatory currents originating more distally [60]. While such a gating effect has its origin in the complexities of neural membrane dynamics, we model it phenomenologically. Thus, let *x*_BG_, *x*_Ctx_, be the level of basal ganglia and cortical inputs, respectively, to thalamus, and *w*_BG_, *w*_Ctx_ the associated synaptic weights. Then inhibition from basal ganglia can act in a gating manner on cortical input if the net input, *u*, to thalamus is given by

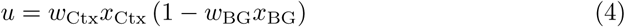

However, projections from SNr to the thalamus synapse both proximally *and* distally [59] on the dendritic arbor, and so we might expect a mix of gating and additive effects. Therefore, we also incorporate additive connections to thalamus from basal ganglia.

The superior colliculus (SC) is a sub-cortical nucleus which also plays a critical role in the generation of saccades [61]. Both FEF and SC have direct connections to the saccadic generator (SG) and if either is lesioned the other can direct gaze following a period of adjustment [62], albeit with some persistent deficits. The SC is also a direct target of output from the SNr [63,64], and thus can be influenced by the action selection mechanisms of the basal ganglia. In particular, it forms a loop with basal ganglia but, unlike its cortical counterpart in FEF, the input to basal ganglia goes via the thalamus.

While the SC has seven alternating cell and fibre rich layers [65], in most cases these are divided into the ‘superficial’ and ‘deep’ layers, which have significantly different response properties. Cells in the superficial layers are mainly visually responsive, with a preferred response to phasic events (luminance onsets and offsets) and movement on the visual field [66], and receive input from the retina. In contrast, cells in the deep layers receive multimodal input, including inhibitory input from the output structures of the BG [63], and are directly involved in the generation of saccadic eye movements. Saccade related activity in the deep layers appears to generate saccades through ‘population coding’, with a weighted sum of activity across the retinatopy of SC determining the saccade target [67–69]. The deep layers of SC receive input from the FEF in a topographic manner [70,71]

We base the SC in our architecture on the model proposed by Arai et al [72]. In their model influence on the SC from the SNr was implemented as a global, rather than ‘channel-based’ signal, that projected to all regions of SC with equal strength; selection of saccade target was mediated by long range inhibitory mechanisms in SC rather than SNr. In a more recent version of the model Arai and Keller (2005) removed the intrinsic inhibitory connections from SC and - in response to new evidence - modelled SNr input to the SC with a coarse topography, thereby allowing SNr to shape selection at a given location. However, unlike our model, Arai and Keller hand-crafted the activity in their SNr input to test a specific hypothesis regarding the role of multiple locus selection in explaining saccade curvature. The Arai and Keller (2005) model of SC has superficial and deep layers which we incorporate here. Each of these layers in our model is a two dimensional array of 50 by 50 leaky integrator units, and as in the FEF is arranged in a retinatopic manner [65].

Consideration of the effect of fixation activity on the model, uncovers a computational problem. Due to its large extent, the foveal region provides a strong salience signal to the BG which could produce sufficient disinhibition to cause saccades. While a peripheral target may ultimately compete successfully with the fovea and generate a saccade, this process would cause a significant delay in producing this saccade, leading to latencies much larger than those observed experimentally. In order to prevent such erroneous behaviour, we cannot simply prevent saccades close to the fovea, as this still allows strong thalamo-cortical loop activity in the foveal representation, which could potentially still be problematic.

A clue to how to proceed is supplied by data showing constant fixation activity in the FEF [22]. This implies that it may not be cortical activity at the fovea which is of concern, but rather, its facilitatory influence downstream. Thus, we suggest a mechanism whereby the synaptic strength of the connections between the FEF, and thalamus and striatum, are reduced closer to the fovea. This is done by multiplying the cortico-fugal weights by a factor *k*(*r*), where *r* is the radial distance from the fovea in pixels (see the following section for details). Thus,

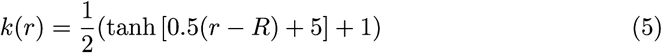

where, *R* is the radius of the neural layer (in pixels). The relation is ‘S-shaped’ normalised to the range [0 1].

### 2.4 Input and output: retina and saccade generation

Input to the model is provided through a simple retina model which directly samples a larger ‘world array’ of pixel values. This input goes directly to FEF via a 50 ms delay, intended to model the passage time through the dorsal visual stream. In this way, FEF provides a sustained response to long lasting visual stimuli. Regions such as the LIP are compacted into this delay, as evidence suggests that while FEF activity is strongly correlated with saccade decisions, LIP activity is not. We therefore assume that accumulation of activity in LIP is a result of the feedback from FEF, where the decision is actually formed. We must also address the lack of motion and other processing in the FEF input. These are omitted as the information is processed in regions which are not on the direct path from the retina to the FEF, and as such will incur greater delays to the information arriving at the FEF. Since decisions are made with low latencies in reflexive saccades we therefore omit this information and simply use the luminance information which can be transmitted most efficiently to the FEF.

In contrast, the SC has a low latency, phasic, response to luminance onset and offset [65]. To capture this phenomenon, we introduce a pair of 50-by-50 leaky integrator unit layers with different membrane time constants, and with the more slowly reacting layer inhibiting its faster counterpart. Both layers take input from the ‘world array’, and the faster layer responds quickly to the appearance of a prolonged stimulus before it is inhibited by the slow layer, forming a phasic response to the stimulus onset; this is then sent to superficial layer of the SC. It should be noted that we do not assert that this is an entirely separate pathway from the retina to the SC, but simply that tonic responses are not found in the SC superior layers, and it is this feature we seek to reproduce.

The retina processes images from a Cartesian ‘world array’ (Fig 3) and, in doing so, captures the nonlinear topographic mapping from image space to its representation in areas like SC and FEF, in which the fovea (central region around point of gaze) is represented much more densely than the periphery. This is due to inhomogeneity of representation in retina and its subsequent projections to target areas like cortex [74]. However we subsume all such transformations using a single cortical magnification factor (CMF) defined by a function *M* (*E*) of the visual eccentricity *E*. We adopt the form for *M* (*E*) in [75]

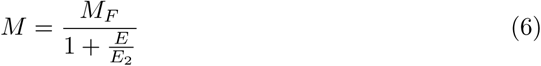

Where *M*_*F*_ the foveal magnification, and *E*_2_ is the eccentricity at which the magnification factor has changed by a factor of 2. There are a range of experimentally measured values for *E*_2_, mostly between 0.1° and 4° [76]; we chose a value of 2.5°.

We now turn to the motor output; saccade generation. While there are several biologically realistic neural models of the saccadic generator (SG), their use would require the crafting of appropriate weights from the topographic array of SC neurons to SG. This in turn requires a learning phase which, given the time required to compute a single saccade in the model, would be prohibitively lengthy. Instead, we make use of a simple interpretation of the retinatopic arrangement of SC to determine the putative saccade position. Here, we first transform the retinatopic organisation of the SC to Cartesian co-ordinates, and then compute the centroid of the activity of the SC deep layers. The position of this centroid in the Cartesian frame determines the saccadic position. This mechanism has previously been used in the target acquisition model of [77].

The production of clean saccades, free from artefacts, is dependent on precise signal timing. In particular, it depends critically on the cessation or ‘quenching’ of target representations after a saccade has been made. Thus, suppose the target representation in FEF is not quenched rapidly enough after a saccade towards the spatial location it originally indicated. The residual activity will indicate a new, spurious target position, bearing the same relation to the new foveal representation as its previous counterpart. This activity, if expressed strongly enough, has the potential to cause a saccade to the new ‘phantom target’, as it were. In extremis, the complete failure to reduce the target activity in FEF can result in a rapid succession of so-called ‘staircase saccades’. This is not simply a theoretical construct; it can be induced by electrical stimulation of the SC, and is found as a symptom in Parkinson’s patients [78–81].

To correct this pathology requires a post-saccadic inhibition of the target location. Such a ‘reset’ signal is available via re-entrant (i.e. feedback) connections from the brainstem to the FEF and SC, ’ [82]. Experimental evidence shows the effect of this signal is suppression following visual activity that evokes a saccade, but not following sub-saccade threshold visual activity.

In addition we suppress all visual input from the target during the saccade, to model the phenomenon of ‘saccadic suppression’ [83]. Since we do not model the mechanics of the eyeball, our saccades are nominally of zero duration (we are not interested in the current model in the dynamic profile of saccades themselves, but rather, their initiation). However, we suppress visual input for 10ms to mimic the finite duration of visual suppression accompanying a saccade.

### 2.5 Model parameterisation and detail

Now we describe in detail the parameters set in the model. These are determined by tuning the model to perform a simple saccadic task, in which a fixed luminance point is fixated by the model. After a fixed duration (since our model cannot learn the timing) the fixation point is extinguished, and simultaneously a target point of fixed luminance is presented. The *saccadic latency* between presentation of the target and the initiation of an eye movement is measured and averaged. We tune the model to match this to experimental data, while also matching the electrophysiological evidence of activity in a variety of brain regions. More details are provided in the Results section. It should be noted that the tuning of the BG portion of the model attempts to preserve as closely as possible the weights used in the original paper.

Fig 3 shows the detailed model structure highlighting several of the features described above (and not apparent in the macro-scale diagram of Fig 1). Projection schemes are one of several varieties: Gaussian spread of weights (see section: Basal ganglia), one-to-one, all-to-all, or normalised in which the sum of all weights in the connection is one. As detailed in the previous section, for the input to the Striatum the weights diminish gradually towards the fovea. All layers have noise applied to the activation level of each neuron (Gaussian, *σ* = 0.01) at each time step of the simulation, except the retinal and world layers.

The parameters for each layer are shown in Table 1

**Table 1.**
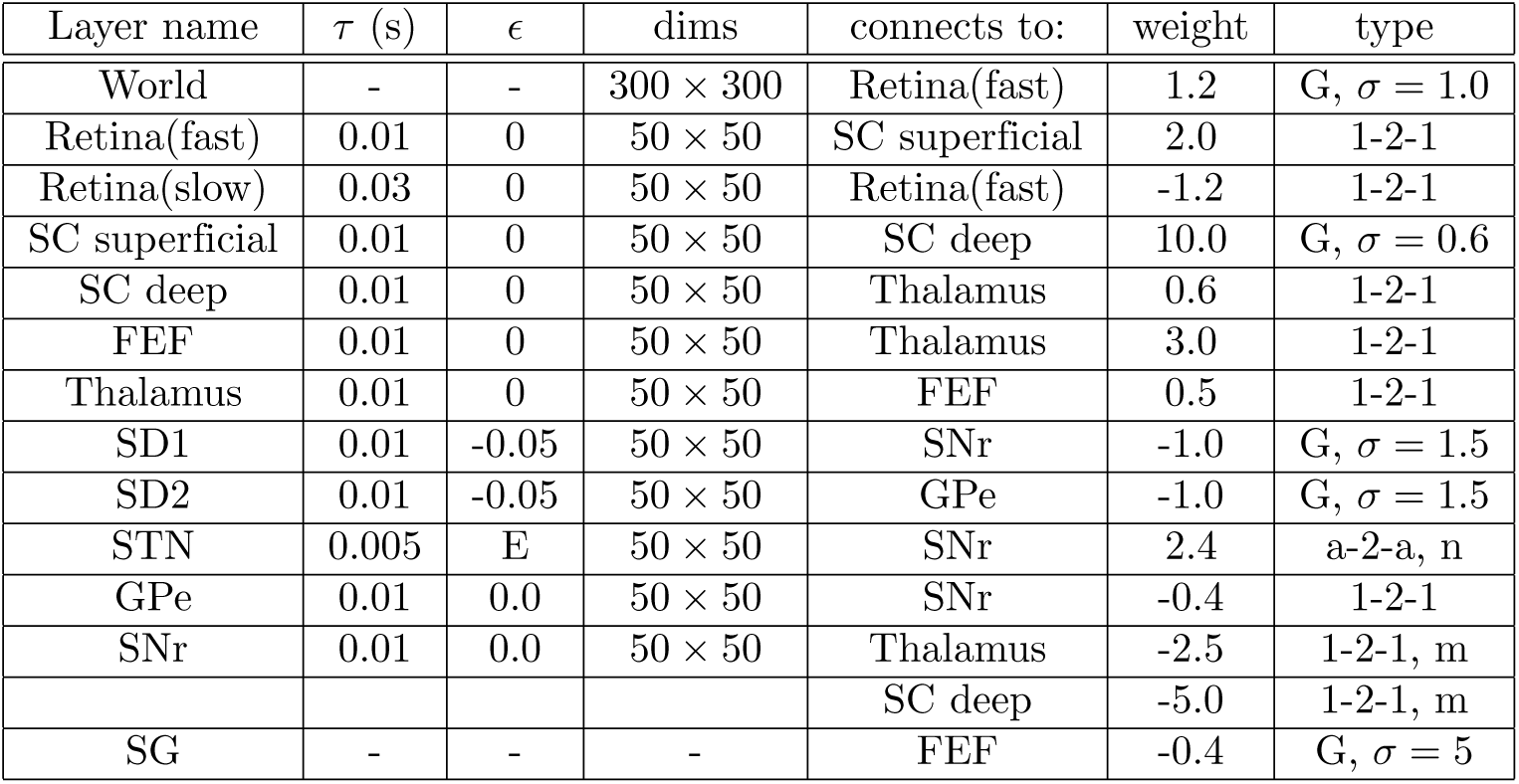
Model parameters: connection strengths, connectivity type, and membrane time constants *τ*. Abbreviations: G; Gaussian projection connectivity with standard deviation of weights *σ*. 1-2-1, a-2-a; one to one, all-to-all connectivity respectively. n: normalised connectivity (see text). df: connectivity grows weaker at the fovea see equation (5). m: multiplicative synaptic. E: exponential output function (see text) links described in (4).

| Layer name | $\tau$ (s) | $\epsilon$ | dims | connects to: | weight | type |
| --- | --- | --- | --- | --- | --- | --- |
| World | - | - | $300 \times 300$ | Retina(fast) | 1.2 | G, $\sigma = 1.0$ |
| Retina(fast) | 0.01 | 0 | $50 \times 50$ | SC superficial | 2.0 | 1-2-1 |
| Retina(slow) | 0.03 | 0 | $50 \times 50$ | Retina(fast) | -1.2 | 1-2-1 |
| SC superficial | 0.01 | 0 | $50 \times 50$ | SC deep | 10.0 | G, $\sigma = 0.6$ |
| SC deep | 0.01 | 0 | $50 \times 50$ | Thalamus | 0.6 | 1-2-1 |
| FEF | 0.01 | 0 | $50 \times 50$ | Thalamus | 3.0 | 1-2-1 |
| Thalamus | 0.01 | 0 | $50 \times 50$ | FEF | 0.5 | 1-2-1 |
| SD1 | 0.01 | -0.05 | $50 \times 50$ | SNr | -1.0 | G, $\sigma = 1.5$ |
| SD2 | 0.01 | -0.05 | $50 \times 50$ | GPe | -1.0 | G, $\sigma = 1.5$ |
| STN | 0.005 | E | $50 \times 50$ | SNr | 2.4 | a-2-a, n |
| GPe | 0.01 | 0.0 | $50 \times 50$ | SNr | -0.4 | 1-2-1 |
| SNr | 0.01 | 0.0 | $50 \times 50$ | Thalamus | -2.5 | 1-2-1, m |
|  |  |  |  | SC deep | -5.0 | 1-2-1, m |
| SG | - | - | - | FEF | -0.4 | G, $\sigma = 5$ |

## 3 Results

One of our aims is to discover the mechanistic explanation of a range of reflexive saccadic *behaviours*, together with the *neural responses* found in the oculomotor system during saccade production. To this end, we investigated several kinds of stimulus paradigm, all of which make use of simple luminance stimuli with limited spatial extent (typically small disks of light) [1,3]. Trials are usually commenced with a *fixation point* and, after a period, the fixation point is extinguished and another point (the *target point* is presented a different location. The time taken from presentation to saccade is the *response time* (RT) or the *saccadic latency*. The paradigms we model look at the several dimensions along which such protocols can be constructed. Some protocols examine the effect of temporal variations in fixation offset and target onset: if the target appears before, simultaneously with, or after, the target offset, we produce so-called *overlap*, *step*, or *gap* stimuli [4]. Other protocols vary the spatial relationship between fixation and target in terms of the visual angle or *eccentri*city between the two [5]. In experiments critical to our aim of examining the role of basal ganglia in action selection, we examine the ability to select between two, simultaneously presented targets [6]. Finally, as well as recording trends of mean RT against independent stimulus variables (like stimulus gap/overlap or eccentricity), much previous research has been directed at uncovering the statistical distribution of RTs for a given parametric setting [3].

### 3.1 Parameterisation: the model can be tuned to account for neural responses during a saccade task

The task was a step-protocol with an isoluminant point fixation (luminance = 0.5 (on a scale from zero to one) and point target (luminance = 0.6, duration 250ms) presented at a fixed retinal eccentricity of eight degrees. The responses of several units in the model were recorded and compared with neural responses recorded from a monkey performing the same task (Fig 4) in order to tune the model parameters. All panels of Fig 4 display an 800ms time window so the duration of changes can be compared across conditions and nuclei. Note that no scaling of simulation time was made to match events in the data; the temporal scale of the dynamics emerges naturally from the model itself.

**Fig 4.**
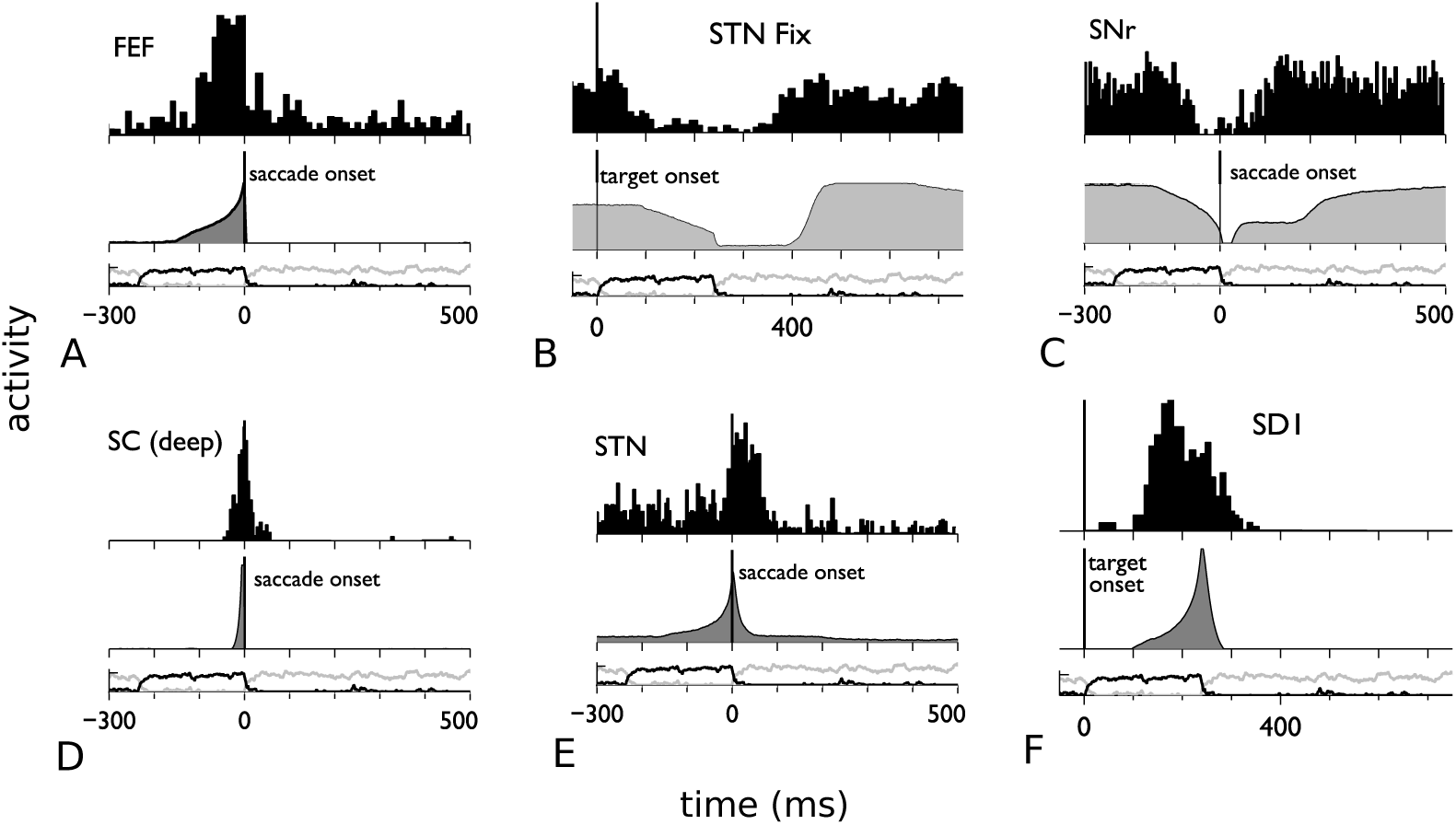
Data and model comparison at the level of neural responses. In each panel, peristimulus time histograms (PSTHs) of neural responses in key oculomotor nuclei are compared with comparable model outputs for a step task; the PSTHs are at the top (black area fill) and the model output below (grey area fill). At the bottom of each panel is shown the retinal input at fixation and target as grey and black line plots respectively. All neurons (model and data) have receptive fields (RFs) covering the target location apart from that in B, (‘STN fix’) which is for STN neurons with RFs encompassing the fixation point. The timings are aligned to saccade or target onset as indicated in each panel and all panels show an 800ms time window. Data were adapted from the following sources: SNr and SC [61]; STN [84]); striatum SD1 [85]); FEF [86]

The results show that the model can produce similar phasic activity increases related to the target stimulus onset in the FEF, SC, STN, and striatum when performing the same task. These increases are associated with a phasic decrease in SNr activity in cells whose receptive field contains the target. Activity in the STN at fixation shows an initial period of high activity due to the fixation stimulus, then a period of low activity following the saccade, followed by high activity due to the relocation of the target to fixation.

We now seek to explain these results, and thereby their biological counterparts, in terms of mechanisms available within the model. Each channel corresponds to a retinatopic location, and a local luminance stimulus will activate one or more of these channels, depending on the retinal area it occupies. Just after target onset, the superficial layer of SC has phasic response to this event, while the FEF starts to respond to the persistent presence of this stimulus – see Fig 4A.

Both FEF and SC provide input to the basal ganglia: the FEF directly, and the SC superficial layer through a relay in the thalamus. Thus, stimulus activity in FEF and SC provides retinatopically organised excitatory input to the striatum and the STN. These inputs start to induce response in STN and SD1 (Fig 4E,F). Note that STN activity at the target location initially shows a non-zero tonic response which, in the model, is due to background noise from the FEF; there is no corresponding activity in the SD1 neurons which are in the down-state). Then, under the off-centre on-surround circuit described in section 2.2, SD1 starts to *decrease* inhibitory SNr activity for the target-location channels (4C), while STN causes *increased* SNr output on all others.

At the level of the loops with FEF and SC, the SNr sends its inhibition to the thalamus and the deep layer of the SC so that, decreasing activity on a channel in SNr disinhibits the corresponding channel in these layers. The deep-layer SC units are more strongly inhibited than their thalamic counterparts, so the latter responds first. The thalamus is in an excitatory loop with the FEF, and so disinhibition from the SNr allows activity in corresponding channels the FEF to increase. This, in turn increases the input on those channels to the basal ganglia resulting in further disinhibition on the selected channels. This feedback ‘amplification’ process is evident in the accelerated rate of change of the signals in FEF, STN, SD1 and SNr (4A,E,F,C). This process continues until the strong inhibition to the SC deep layer is almost completely removed, allowing a vigorous phasic response in this layer (4D) which initiates a saccade.

Once the saccade has been initiated, the visual input changes, as the saccade has moved the target to the centre of vision. There is an accompanying rapid decay of activity in the STN at the previous location of the target on the visual field.

At the foveal fixation point, prior to the saccade, STN receives sustained input from FEF until the fixation is removed, thereby causing a phasic decrease in STN activity at this location (4B). During the saccade, the post-saccadic inhibitory mechanisms remove afferent drive from STN causing its activity to decline. After a delay corresponding to that in the dorsal stream to FEF, in addition to the saccade duration, the retinal activity derived from the target at its new (foveal) location occurs, increasing the activity in STN at that new foveal location. This, in turn, drives SNr activity back up to fixation levels.

### 3.2 Validation: the model accounts for the characteristic pattern of response times for gap, step, and overlap tasks

We now begin the process of validating the model by testing a variety of behavioural experiments that the model has not been tuned explicitly to reproduce. The gap, step and overlap tasks may be viewed as categories of a single task protocol with variable temporal gap between fixation and target stimuli [4]. Thus, we conducted experiments to measure the saccadic reaction time (RT) as we varied the temporal separation, *δt*, between fixation offset and target onset over a range of values between -400ms and 400ms. Here, negative and positive values signal overlap and gap tasks respectively, and *δt* = 0 is a step task (as in the previous section). Luminance and target eccentricity were constant, as was the duration of the target, and all were the same as those used in the previous section.

The results, together with data from [4], are shown in Fig 5. Note the three experimental subjects show considerable individual differences in the precise values of RT, but not on the patterning of the data: there is always a fairly constant set of values for large negative *δt*, linearly decreasing values across the step condition (*δt* = 0), and another region of approximate constancy for large positive *δt*. In respect of these features, the model results show a similar patterning to the data.

**Fig 5.**
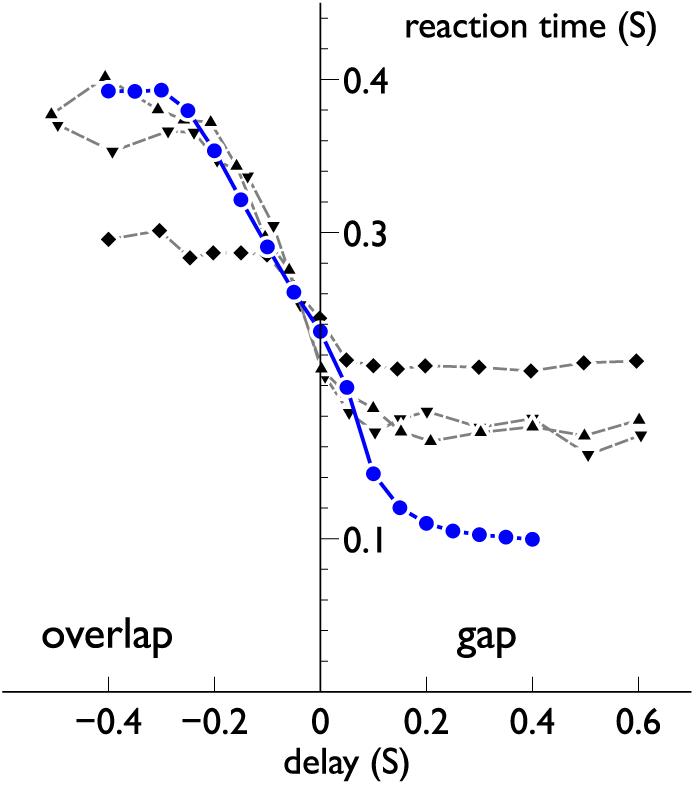
The effect of delay between fixation point offset and target onset on saccadic reaction time. The blue filled symbols and solid line show model results. The three plots with black filled symbols and dashed lines are data from experiments by [4] with three different subjects. Error bars are suppressed for clarity

In order to explain this pattern of results, we examined the behaviour of several neuron types in the model for a gap of 300ms, a step, and an overlap of 300ms (Fig 6)

**Fig 6.**
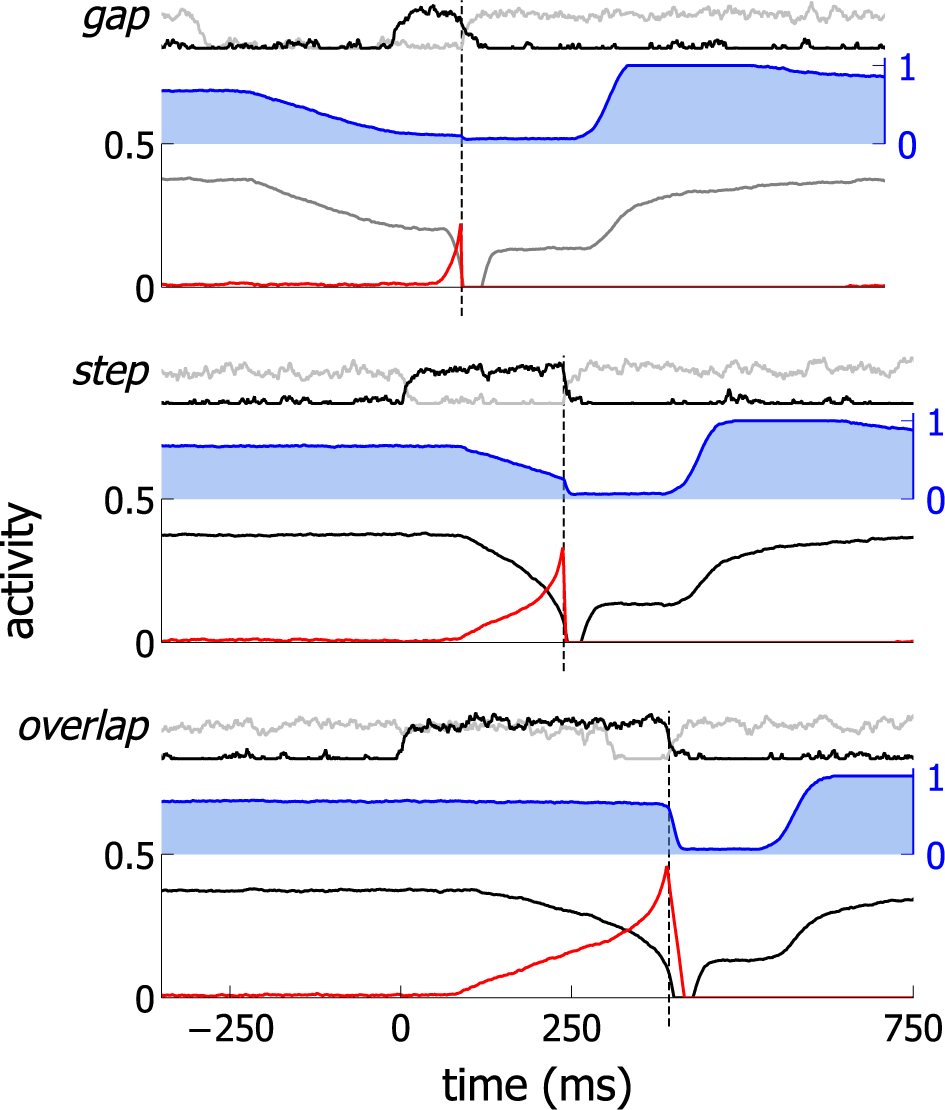
Neural behaviour in the model for gap, step and overlap conditions. Each condition contains subplots as follows: top – retinal input at fixation and target (grey and black line plots respectively); middle – STN activity at fixation ((blue line and shading); bottom – SNr and FEF activity at the target (black and red lines respectively). Graphs are aligned to target onset. The black, dashed vertical lines denote saccade onset.

The three plots are aligned at the target stimulus onset (*t* = 0), and the saccade onset occurs progressively further from this point in going from gap, to step, to overlap conditions. The key to understanding the reason for this is the STN activity. Recall that STN projections are diffuse so that STN activity originating anywhere in the retinatopic array will make itself felt everywhere else in this array. In, particular, activity in STN at fixation influences SNr activity at the target.

In the gap condition, activity in the STN starts to decrease as soon as fixation offsets (and before target onset). This reduces the excitatory signal to the SNr, thereby lowering the level of SNr output. By the time the target point is presented, SNr output has almost returned to its tonic level with no stimulus. The inhibition on the cortico-thalamic loop between the FEF and thalamus is, therefore, comparatively low when the target occurs. This allows the activity in the FEF-thalamic loop to accumulate rapidly, leading to a short RT.

In the step condition, activity at fixation in the STN does not begin to decay until a short time after the target is presented. The decrease in SNr output is therefore correspondingly delayed compared to the gap condition, leading to slower integration in the cortico-thalamic loop, and a longer RT.

In the overlap condition, the fixation activity in STN remains high at the target location for some time after target onset, resulting in SNr activity at the target location taking longer to decrease. This, in turn, causes a longer time for integration in the cortico-thalamic loop and thereby, longer RTs.

A similar reasoning may be used to explain the increased occurrence of very short latency, *express saccades* in the gap condition [3]. Thus, the relatively low inhibition at target onset from the SNr to the deep layer of SC allows activity induced in the SC superficial layer by target onset to stimulate a saccade without the persistent activity from the FEF.

### 3.3 Validation: the model accounts for patterns of data with varying target eccentricity and luminance

In [5], Kalesnykas and Hallett showed how, for the step paradigm, RT depended on the eccentricity of the target (in relation to fixation) for a range of target luminance values. We replicated this kind of experiment using eccentricities from 3 to 40 degrees. In addition the same task was performed with two target stimulus luminances of 0.5 and 0.9 (on a normalised scale).

Fig 7 shows the model results, and data from the experiment with human subjects from [5]. The model results show the main features for the dependency of RT on eccentricity apparent in the data: a ‘hockey-stick’ shape with the knee of the curve at small eccentricities. In addition the decrease in RT with luminance is also shown in the model outcome, along with the preservation of the ‘hockey-stick’ trend across luminance values.

**Fig 7.**
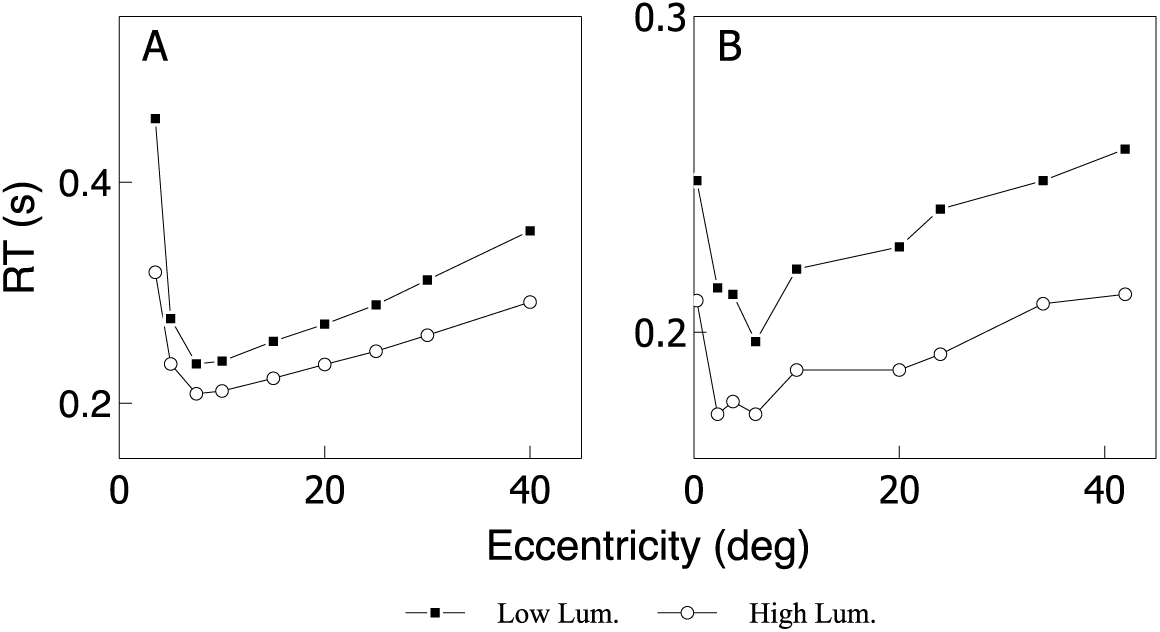
RT as a function of eccentricity and luminance. A, model results for target stimuli with luminances of 0.5 (‘Low’) and 0.9 (‘High’) on a normalised scale. B, adapted from data in [5] showing luminance 10 times and 1000 times greater than the dark-adapted foveal threshold (FT) of the subject (‘Low’ and ‘High’ luminance respectively). In both panels, the High and Low luminance values are shown by open and closed symbols respectively.

In order to explain this behaviour we examine several neural responses at a critical stage in developing the saccade (Fig 8).

**Fig 8.**
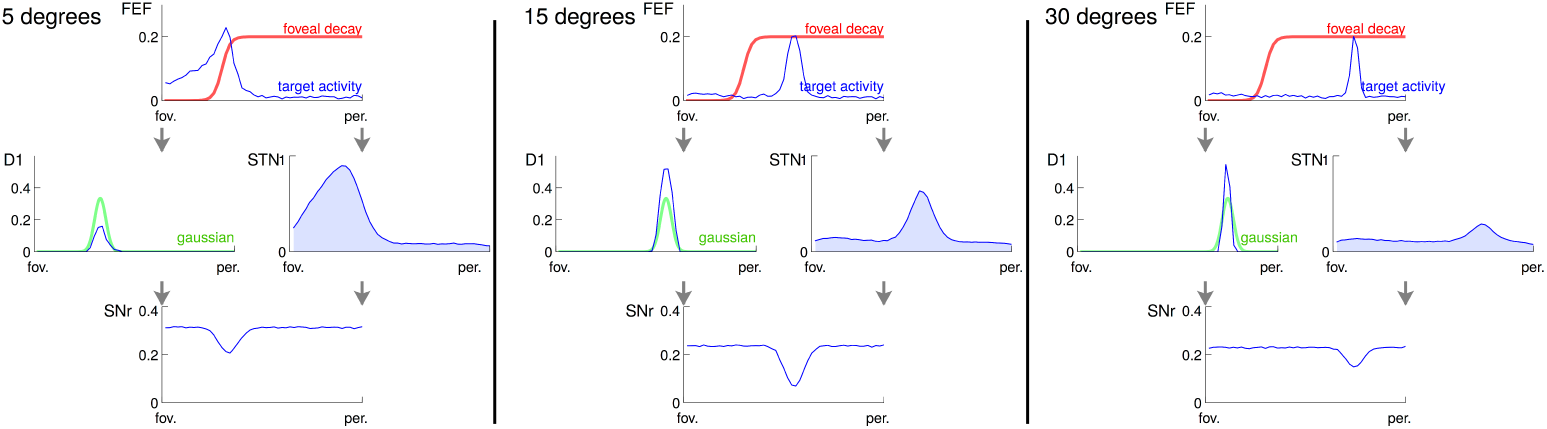
Results from the model for a single time step. This time step is chosen when the FEF activity reaches 0.2. The D2 striatal pathway is not shown for clarity. For each condition (5, 15 and 30 degree target) a cartoon of the activity in the FEF activity, striatum activity and output (top and bottom respectively), STN activity and output (top and bottom respectively), and SNr activity are shown. The effect of the reduction in weight at the fovea for striatal inputs (red), the Gaussian projective field of the striato-fugal projection (green) and summation of STN activity (blue shading) on the target activity, and consequently the SNr output are highlighted. See the text for a full explanation. fov. and per. denote the fovea and periphery of the visual field respectively.

The behaviour as eccentricity increases can then be explained in the model by considering the competition between two loci of activity generated by the target. The first of these is that in the striatum, which will tend to cause release of inhibition at SNr via the striato-fugal projection from SD1, thereby promoting integration in the cortico-thalamic loop with FEF. The second locus of activity is that in STN, which will tend to prevent selection by increasing activity across all channels in the SNr.

We now make a semiquantitative analysis of the contribution of each of STN and SD1, by examining the way in which ‘hills’ of activity in each nucleus interact with a topographically coincident receptive field (RF) in SNr (Fig 9).

**Fig 9.**
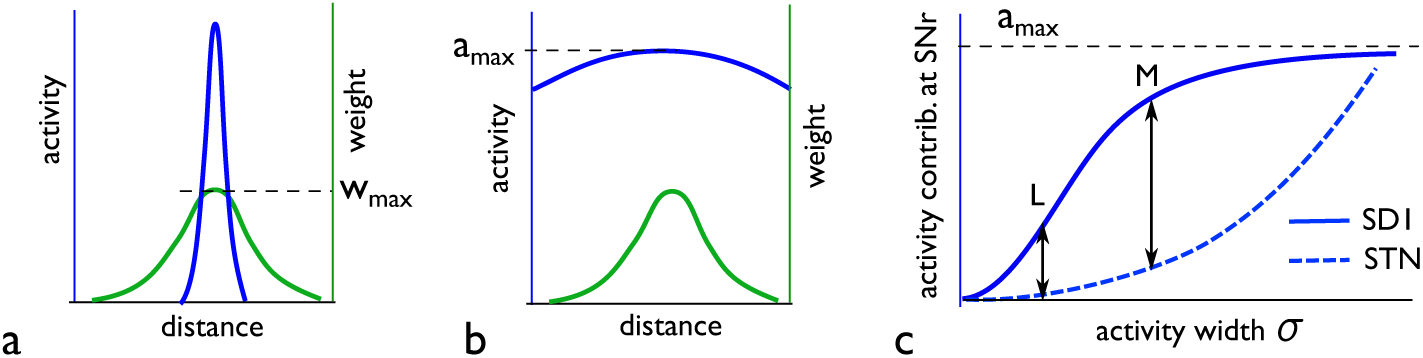
Contributions to activity in SNr from SD1 and STN in the experiments of Fig 8. a, 1D cartoon (blue line) of narrow activity profile in SD1 (a 2D ‘hill’ of activity in the model) as a function of distance across its topographic map. Also shown (in green) is the fixed weight profile for the SNr receptive field from SD1; *w*_max_ is the maximum weight in this RF. b, similar to a, but with a much wider activity profile, only part of which is shown. c, semiquantitative cartoon diagram of the relative contributions to the SNR RF from SD1 (solid line) and STN (dashed line) as the width of the activity profile (governed by a parameter *σ*) in each nucleus increases (see text). The points marked ‘L’ and ‘M’ indicate values of *σ* originating from experiments with ‘large’ and ‘moderate’ eccentricities respectively.

To proceed, assume the activity profiles are isotropic 2D Gaussian functions with width parameter *σ* (which is often a reasonably good approximation). Fig 9a shows a 1D representation of an activity ‘hill’ in SD1 with width *σ*_*n*_ that ensures the profile is much narrower than the RF weight distribution. For such a narrow profile, the weights don’t change significantly over its entire extent, and so we make the approximation that the activity is uniformly weighted by the maximum weight *w*_max_ of the RF in SNr. The resulting contribution *a*_SD1_(*σ*_*n*_) to SNr activity will then be proportional to the volume under the Gaussian: 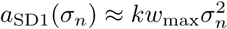. Thus, for small *σ*, *a*_SD1_(*σ*_*n*_) is approximately quadratic in *σ* with slope *kw*_max_*σ*.

Now consider a much wider SD1 activity profile, with width *σ*_*w*_, shown in Fig 9b. However, we assume this has the same peak value, *a*_max_, as its narrow counterpart in Fig 9a. For the wider profile we assume that the activity is approximately constant over the RF, so that the contribution *a*_SD1_(*σ*_*w*_) to SNr activity is given by *a*_SD1_(*σ*_*w*_) = *a*_max_ ∑_*i*_ *w*_*i*_, where the RF is comprised of weights *w*_*i*_. However, SD1 projections are normalised so that *a*_SD1_(*σ*_*w*_) *≈ a*_max_. From the figure it is clear that *a*_SD1_(*σ*_*w*_) *> a*_SD1_(*σ*_*n*_), and so *a*_max_ is an upper bound for *a*_SD1_(*σ*). Bringing these observations together (for small and large *σ*), Fig 9c shows a cartoon of *a*_SD1_(*σ*) (solid line).

We now turn to the contribution from STN which projects diffusely and uniformly to SNr with weight *w*_STN_. The contribution to SNr activity, *a*_STN_, is therefore given by *a*_STN_(*σ*) = *k*′*w*_STN_*σ*^2^, where *σ* is the width of the STN hill of activity and *k*′ plays the same role as *k* for SD1; if the STN peak activity peak is the same as that in SD1, then *k*′ = *k*. Thus *a*_STN_(*σ*) is also quadratic in *σ* and, for small values of its argument, it has slope *k*′*w*_STN_*σ*.

The preceding analyses may now be combined to compare the contributions *a*_STN_(*σ*) from STN, and *a*_SD1_(*σ*) from SD1 to SNr activity. While the sum of the weights projecting to an SNr neuron is 2.4 and that for SD1 projection is 1, the much larger RF with respect to the diffuse projection from STN means that *w*_STN_ ≪ *w*_max_. Thus, unless *k* ≪ *k*′, then for small *σ* where the quadratic approximation for SD1 holds, the slope of *a*_SD1_(*σ*) is much less than its STN counterpart. Further, while the slope of *a*_STN_(*σ*) is monotonically increasing, *a*_STN_(*σ*) itself will only reach the level of *a*_SD1_(*σ*) beyond the quadratic regime (see Fig 9c) so for a significant range of *σ*, *a*_STN_(*σ*) *< a*_SD1_(*σ*).

Returning to Fig 8, consider the activity profiles for 30 degrees eccentricity. The activity in FEF is transmitted faithfully to the BG as the cortico-striatal weights do not diminish (due to the foveal reduction) at this distance. However, the activity in FEF occupies a rather small part of the neural layer as the input is topographically arranged with substantially more resources at the fovea. This results in a correspondingly narrow zone of activity in striatum and so, with respect to Fig 9c, *σ* is relatively small; we assume it occupies point typified by ‘L’ therein.

Now consider the case of 15 degrees eccentricity in Fig 8. The (faithfully transmitted) FEF activity now occupies a larger area of the neural layer than it did for the case of 30 degrees, resulting in a wider ‘hill’ of activity in striatum and STN. Thus, in Fig 8, *σ* is greater than it was at the eccentricity of 30 degrees and we assume it occupies a point typified by ‘M’ therein. The relative influence of SD1 compared to STN is larger here than it is at the point ‘L’ (indicative of the 30 degree eccentricity), which accounts for the increased inhibition at SNr. This, in turn, causes a faster rate of accumulation in the cortico-thalamic loop leading to a smaller RT (see Fig 7).

When the target is at the 5 degree eccentricity, the horizontal extent of the hill of target-related activity in FEF is wider still. Thus, in Fig 9c, a corresponding *σ* would be to the right of its value at ‘M’. This may result in a larger difference between SD1 and STN than that for larger eccentricities (such as 15 degrees) but, this is not guaranteed. However, there is a potentially more profound influence at work here which was not in play for the 15 and 30 degree targets; that is, the activity of the target partially overlaps with the weakened zone of cortico-striatal and thalamo-striatal weights at the fovea (see the red line in Fig 8). This results in a decrease in the Striatal inhibition of SNr (in comparison with the 15 and 30 degree targets), and therefore a corresponding decrease in the rate of accumulation in the cortico-thalamic loop, resulting in an increase in RT at low eccentricity seen in Fig 7.

The reduction in RT due to increasing luminance of the target point seen in Fig 7 is produced in the model by the increased peak height of the persistent luminance signal into the FEF. The higher signal strength means that integration in the cortico-thalamic loop proceeds faster than with a weaker signal, thus leading to disinhibition of the SC and the generation of a saccade in a shorter time from the target onset.

These experiments highlight two main mechanisms in the model which can affect the RT of a saccade. The first is the competition between inhibition and excitation to SNr from SD1 and STN respectively. This, in turn, relies on two factors: (i) a complex interplay between the projective fields from SD1/STN to SNr, and the activity profiles in the former induced by FEF; (ii) the way in which FEF-induced activity in striatum depends on the weight roll-off towards the fovea. The second major mechanism is the strength, or salience, of the target input to the FEF, which leads to increased disinhibition at the target location with increased target luminance or size and thus decreased RT. It is the balance between these opposing mechanisms which creates the ‘hockey-stick’ relationship between eccentricity and RT.

### 3.4 Validation: the model can account for increased ‘saccade averaging’

In the experimental protocol investigated here, the fixation point is extinguished and two targets are presented with equal luminance. They are located an equal distance from the fixation point, but at angular separation *α* and RTs are measured as a function *α* [6]. *The value of α* is varied to 30, 60 and 90 degrees and the timing is such that there is no delay between fixation offset and target onset (a step paradigm). The task is to saccade to either target. There is therefore a competition between the them, posing a genuine action selection problem.

Typically saccades are either to the individual targets or somewhere in-between in which case it is deemed to be an example of *saccade averaging*. More precisely, the metric used to report the results is the percentage of saccades outside the 95% confidence interval of the positions found when single targets were used [6]. Fig 10 shows examples of data from [6] together with the results of the simulation of this experiment using the same metric. There is good agreement between the model and the data.

**Fig 10.**
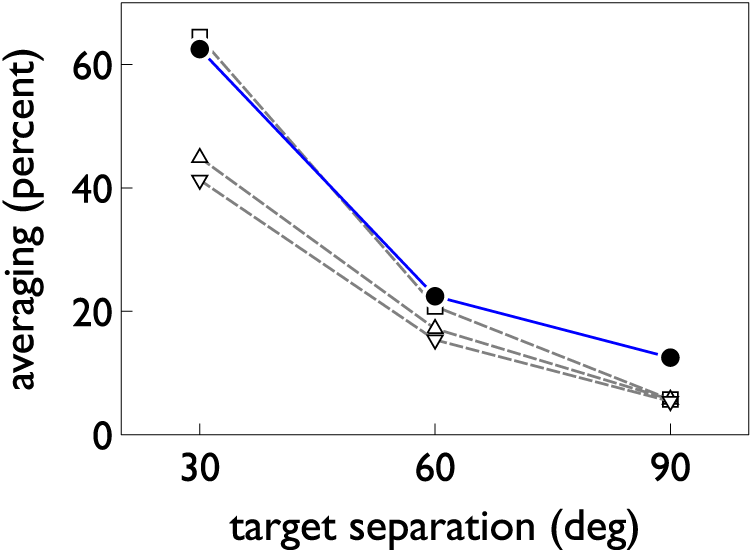
Saccade averaging: data and model. The open symbols and grey dashed lines show the results from three different primate subjects in experiments conducted by Chou et al [6]. The solid symbols and blue line is the result from the model.

An explanation of the phenomenon of saccade averaging is found in the model by invoking the finite width of the receptive fields in the BG. In the thalamocortical loop, this causes both targets to build up ‘hills’ of activity across the retinatopy of their neural layers. As the distance between the targets decreases these hills begin to overlap and interact. The precise outcome of this interaction is noise dependent and results in one of three possibilities: (i) amalgamation of the activity hills for each target into a single hill associated with a location between the two targets; (ii) two distinct hills of activity in which the competition between the two targets has failed to resolve in favour of one of them; (iii) a single hill of activity associated with a single target, arising from complete and successful inter-target competition. Cases (i) and (ii) will give rise to average saccades. Some typical examples are shown in Fig 11.

**Fig 11.**
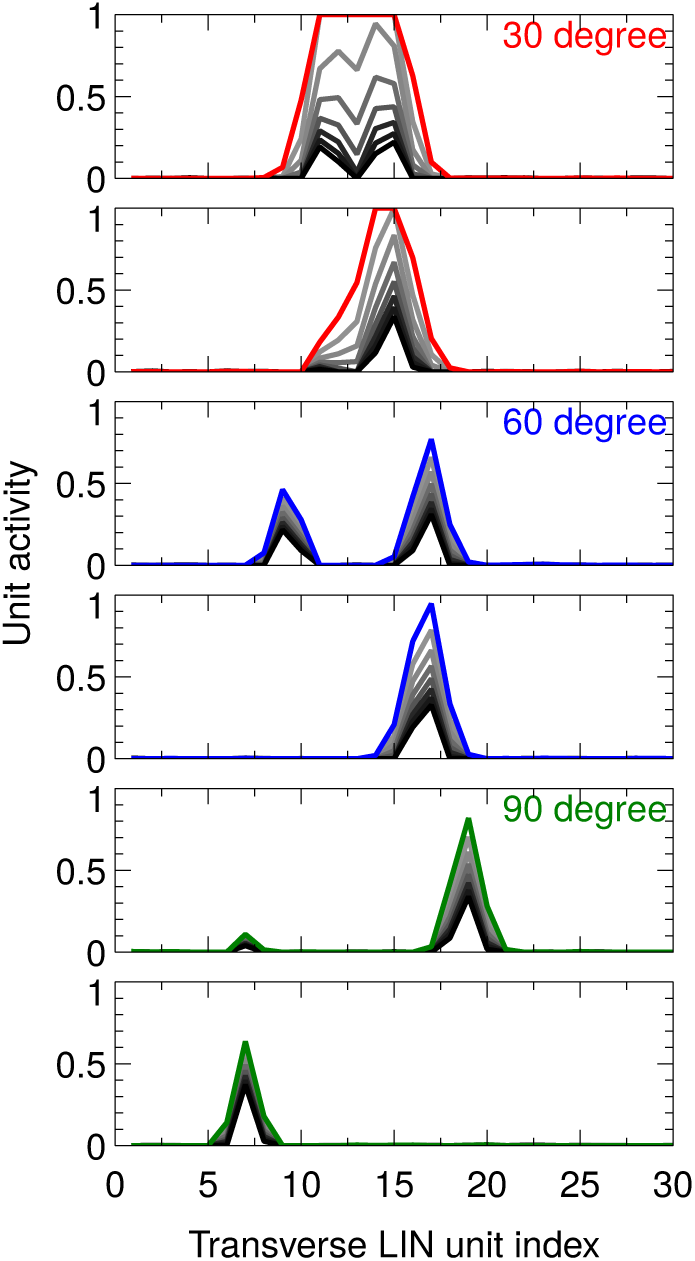
Explanation of saccade averaging. Each panel shows the time course of activity in the deep layer of SC in a saccade averaging experiment. Activity is shown for a set of neurons (indexed along the *x*-axis) along a line in the retinatopic array encompassing the two targets. The heavy black line is an early activity profile, lighter grey lines show snapshots at later times, and the final profile is indicated in colour for each of three target pair separations – 30°, 60°, 90°. For each target separation, two instances are shown. The upper panel of each instance-pair shows the case when the hills of activity become merged (30° only) or competitively unresolved (two hills remain). The lower panel of each pair shows the case when only one hill of activity remains after successful competition between the target stimuli.

For an inter-target distance of 90°, there are few instances of (ii) and, when the do occur the spurious activity hill is very small so that the average position is not very far from that indicated by the dominant hill of activity. For 60°, the consequences of averaging are more substantial, while at 30° this is the main outcome.

### 3.5 Validation: the distribution of saccadic reaction times

The distributions of saccadic RTs are highly variable and can include unimodal distributions, and multimodal distributions exhibiting several peaks [3,87–89]. In reporting these variations, there are two experimental manipulations which often occur. First, the gap, step and overlap paradigms are contrasted, and typically, there is a decrease in RT in the gap paradigm compared with the step and overlap variants [90]. Second, under extended testing over many blocks of trials, subjects often show the ability to increase their performance (smaller RT) over several blocks (especially if they are initially naive to the task) [91]. In both cases the smaller RT is often associated with a new, short latency mode or peak in the response distribution at around 100-130ms. The saccades comprising this peak are termed *express saccades*. Saccades contributing to the peaks at higher RTs are termed *regular saccades*. There is, however, large individual difference between participants. Some show clear multimodal RT distributions [87,89], while others show no discernible multimodality [92,93].

We hypothesised that the separate model loops dealing with phasic and persistent information could represent the different modes of saccadic production; that is, the loops involving SC and FEF enable express and regular saccades, respectively. To investigate this, we performed manipulations to the model to alternately lesion these loops. To lesion the phasic loop we removed the connection from the retina to the SC, and to lesion the tonic loop we remove the connection from the world to the FEF.

The presence of two distinct pathways with characteristic processing times matching the two saccadic modalities could also provide an explanation for the generation of an express saccade peak through training. Thus, in the saccade task, rapid saccades to small phasic onsets are strongly rewarded and, given large blocks of repeated trials, we propose that the neural circuitry will adapt to prioritise the more timely phasic information over the slower and redundant tonic information. We model this effect here by increasing the weight from the model’s phasic processing retinal layer to the superficial layer of SC, thus increasing the strength of phasic input to selection.

We ran experiments with the step and gap paradigms similar to those described earlier; the gap variant had a fixation-target interval of 100ms (step has 0ms). To investigate training, experiments were conducted in two blocks of 800 trials and, between blocks, the weight between the retina and SC-superficial populations was increased from 1.0 to 3.0. The results for all four conditions, showing the distribution of saccades against RT, are shown in Fig 12.

**Fig 12.**
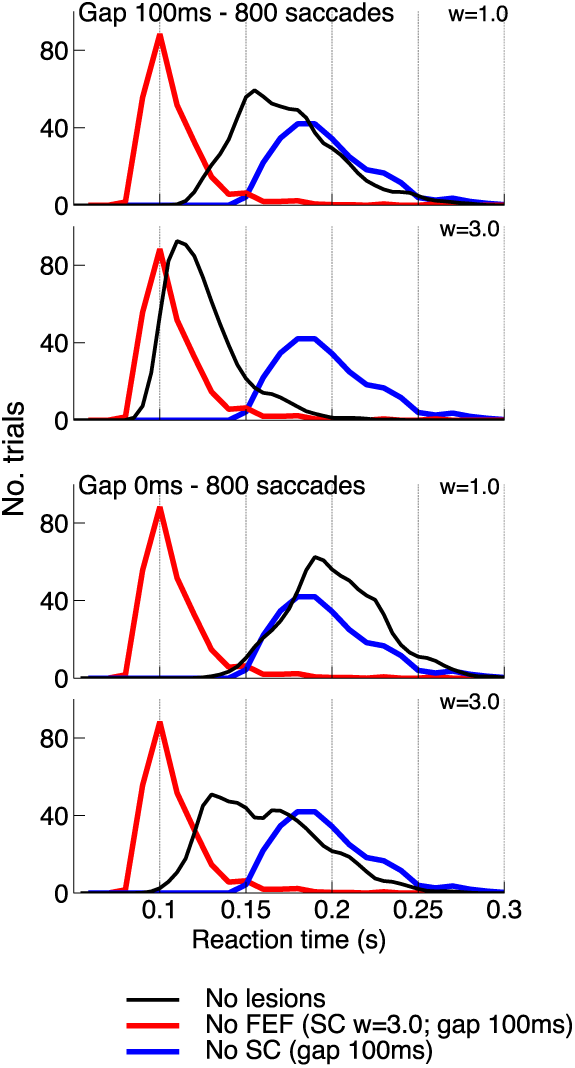
The distribution of saccades generated by the model for several conditions. The red line shows results with lesioned FEF, a gap of 100ms and retina-collicular weight, *w*, of 3.0. The blue line is for a gap of 100ms but no SC. The black line shows the unlesioned results for the conditions indicated in each panel.

There are, as predicted, two distinct modes of performance associated with each lesion. With lesioned FEF, the model used the ‘phasic-only’ loop (involving SC) and gave a peak RT around 100ms, consistent with express saccades. In contrast the ‘tonic-only’ model with lesioned SC (which relied on FEF) gave a peak around 170ms, indicating regular saccades.

For the gap paradigm, increasing the strength of the phasic retinal input leads to a shift in the RT distribution from regular saccades to express saccades. For the step paradigm (gap = 0ms) there is also a shift towards lower RT, however there are much fewer saccades within the express saccade region than with the gap paradigm.

We have shown that the phasic and persistent pathways in the model produce different distributions of saccades that match the RT distributions of express and regular saccades respectively. This agrees with previous explanations of RT distributions that invoked multiple pathways (e.g. [88]). However, in contrast to these earlier models which use abstract functional pathways, our model is grounded in the anatomy and was not constructed to describe express and regular saccades *per se*; these emerge from candidate pathways that occur naturally in the architecture. (There is evidence for a third, slower pathway involving a loop through the prefrontal cortex, or through the ventral stream of the visual system, but this is beyond the scope of our current model).

Despite these distinct pathways, we do not find a multimodal distribution of saccades in the unlesioned model. This is not surprising, however, as 60% of recorded data sets found by Gezeck et al [89] showed no bimodality in their analysis. It is therefore possible that the individual differences shown in experimental subjects can be explained by manipulations to the weights in the model, however such an analysis is beyond the scope of this paper.

The failure to generate many express saccades in the step condition matches experimental findings, and in our model can be explained using the same mechanism invoked to account for the relative increase in RT in the the step condition (compared to the gap condition). After the fixation point is removed, activity related to fixation still exists in the STN, and this activity decreases the rate with which activity in the cortico-thalamic loop can build and generate a saccade. This results in the suppression of express saccades for the step condition.

## 4 Discussion

In this paper we present the integration of an existing biologically plausible model of the Basal Ganglia into a complete sensorimotor system governing saccadic eye movements. Previously such integrated models have been used to investigate voluntary tasks, such as memory guided saccades [13]. With this model we instead focus on investigating the accuracy with which an integrated BG system can reproduce a suite of reflexive saccadic experimental paradigms. The resulting model of the oculomotor system is able to reproduce several observed reflexive saccadic oculomotor behaviours, despite only having been tuned to reproduce simple saccadic step-paradigm behaviour and neurophysiological traces. Moreover, the model is able to provide explanations for these behaviours in terms of the propagation and processing of signals in key brain structures, consistent with electrophysiological data.

The model is fairly complex, with many structures and several feedback loops. However, the explanation of some of the behavioural phenomena (such as the ‘hockey-stick’ for RT against eccentricity) rely on a complex interplay between several mechanism that would not be available in a simpler model. Thus, while we don’t claim that we have accounted for all contributory mechanisms to all examined phenomena, we do argue that some of these phenomena are intrinsically complex, and require a minimal anatomical veracity to account for them properly. These models should then be treated as ‘in silico preparations’ which can be continuously ‘mined’ for explanatory power.

Our model is based on a well established anatomical and neurophysiological architecture, however it is pertinent to compare the explanatory power of our model with that of more abstract models of reflexive saccadic eye movements that can be found in the literature. Influenced by field concepts in physics, the dynamic field theory of movement preparation as described by [94], is a theoretical framework that was devised to explore the generation and modulation of motor signals in the brain. The theory posits that movement parameters are encoded by an activation field that is spatially distributed over some computational space. Under this scheme, both the input supplied to the space, and local dynamic interactions within it, act to shape the evolution of the activation fields and the movements they give rise to. It is no coincidence that this view of action generation maps naturally onto the spatial maps observed within the superior colliculus (and other oculomotor structures).

The models of Klaus Kopecz [95,96] and [97] attempt to represent the saccadic system as a dynamic field model. The most biologically constrained of these is the model of [97] in which the buildup and burst cells of the intermediate SC are modelled as distinct dynamic fields and attempts are made to fit the time course of neural activity observed in primates performing oculomotor tasks. This model provides insights into a number of phenomena affecting saccadic reaction times, including the step, gap and overlap effects and the effect of distractors i.e. multiple simultaneous target stimuli. Furthermore, by making assumptions about endogenous signals supplied to the intermediate SC during voluntary paradigms, the authors are also able to reproduce reaction time effects associated with the anti-saccade paradigm, and with varying target probabilities.

It is impressive that the relatively simple model of [97] and similar dynamic field models, are able to reproduce such a rich set of saccadic phenomena. The computational principles invoked to explain RT effects in such models are the same as those found in the model presented here, namely that spatially distributed population codes that have a tendency to persist, can cooperate or compete depending on their location in their embedding space, and that this competition can affect both the time taken for a movement to be initiated, and its resultant metrics.

Despite this, these models are rather abstract in nature because, in so far as they attempt to capture results that are demonstrably reliant on brain areas outside of the SC, it is clear that they can, at best, only do so at a coarse grained level of description. This contrasts with the model presented in this article, which is heavily constrained by the results of anatomical and physiological experiments, and provides explanations which are wholly tied to the biology. For example, it is difficult to see the results related to saccade reaction time distribution emerging from a simple model of SC, as they do from the model presented here.

While our model accounts for a range of saccadic behaviours and physiological signatures it does not account for the phenomena of voluntary saccades or learning. For this we would need to integrate the competencies shown in existing models [13]. Our model does, however, highlight the continued power of the GPR basal ganglia model, demonstrating the ability of the model, as part of an extended system, to replicate both qualitatively and quantitatively a wide range of reflexive saccadic experimental paradigms. Such validation supports future work in integrating the GPR model as the decision making basis of more complex models of the primate brain.

